# Multimodal Study of Resting-State Functional Connectivity Networks using EEG electrodes position as seed

**DOI:** 10.1101/167585

**Authors:** Gonzalo M. Rojas, Carolina Alvarez, Carlos Montoya, María de la Iglesia-Vayá, Jaime Cisternas, Marcelo Gálvez

## Abstract

Electroencephalography (EEG) is the standard diagnosis method for a wide variety of diseases such as epilepsy, sleep disorders, encephalopathies, and coma, among others. Resting-state functional magnetic resonance (rs-fMRI) is currently a technique used in research in both healthy individuals as well as patients. EEG and fMRI are procedures used to obtain direct and indirect measurements of brain neural activity: EEG measures the electrical activity of the brain using electrodes placed on the scalp, and fMRI detects the changes in blood oxygenation that occur in response to neural activity. EEG has a high temporal resolution and low spatial resolution, while fMRI has high spatial resolution and low temporal resolution. Thus, the combination of EEG with rs-fMRI using different methods could be very useful for research and clinical applications. In this article, we describe and show the results of a new methodology for processing rs-fMRI using seeds positioned according to the 10-10 EEG standard. We analyze the functional connectivity and adjacency matrices obtained using 65 seeds based on 10-10 EEG scheme and 21 seeds based on 10-20 EEG. Connectivity networks are created using each 10-20 EEG seeds and are analyzed by comparisons to the seven networks that have been found in recent studies. The proposed method captures high correlation between contralateral seeds, ipsilateral and contralateral occipital seeds, and some in the frontal lobe.

## 1 Introduction

Electroencephalography (EEG) is a method that measures the electrical activity of the brain. EEG uses surface electrodes to measure the electrical brain signals (Niedermeyer et al., 1993; Tatum et al., 2008).

EEG is routinely used to diagnose or monitor the following medical conditions and diseases: differentiate epileptic seizures, pre-operative assessment for defining an eventually resectable epileptogenic zone for a successful surgery, tumors, stroke, differential diagnosis of paroxysmal events, analysis of encephalopathies, monitoring consciousness, sedated patients at risk of seizures, patients in whom the only way to monitor brain function is EEG (Extracorporeal membrane oxygenation, for instance), prognosis cardiac arrest and hypoxic ischemic encephalopathy (HIE), brain death diagnosis, psychomotor regression study, acoustic or language development, suspected specific genetic phenotypes, and sleep disorders (Bickford, 1987). EEG is used for research purposes in the following topics: neuromarketing, psychology (processes underlying attention, learning and memory).

The International Federation of Clinical Neurophysiology (http://www.icfn.info) adopted the standardization for EEG electrode placement called 10-20 electrode placement protocol (Jasper, 1958; Klem et al., 1999). This protocol standardized the physical placements and designations of 21 electrodes on the scalp. Using reference points on the skull in the nasion, preauricular points and inion (Figure 1), the head is divided into proportional positions to provide adequate coverage of all the brain regions. The 10-20 designation indicates the fact that the electrodes are placed at 10%, 20%, 20%, 20%, 20%, and 10% of the total nasion-to-inion distance along the midline. The other series of electrodes are also placed at similar fractional distances from the corresponding reference distances. The name of each electrode consists of a letter and a number. The letter refers to the region of the brain where the electrode is positioned (F: frontal, C: central, T: temporal, P: posterior, and O: occipital), and the number is related to the cerebral hemisphere (even numbers in the right hemisphere, and odd numbers in the left; Figure 1). In 1985, an extension to the original 10-20 system was proposed involving an increase in the number of electrodes from 21 to 74 (Figure 2) (American Electroencephalographic Society, 1994; Chatrian et al., 1988; Klem et al., 1999; Nuwer et al., 1998). 10-20 EEG electrode placement system is considered for clinical use, and 10-10 is more used for research.

**Figure 1:**
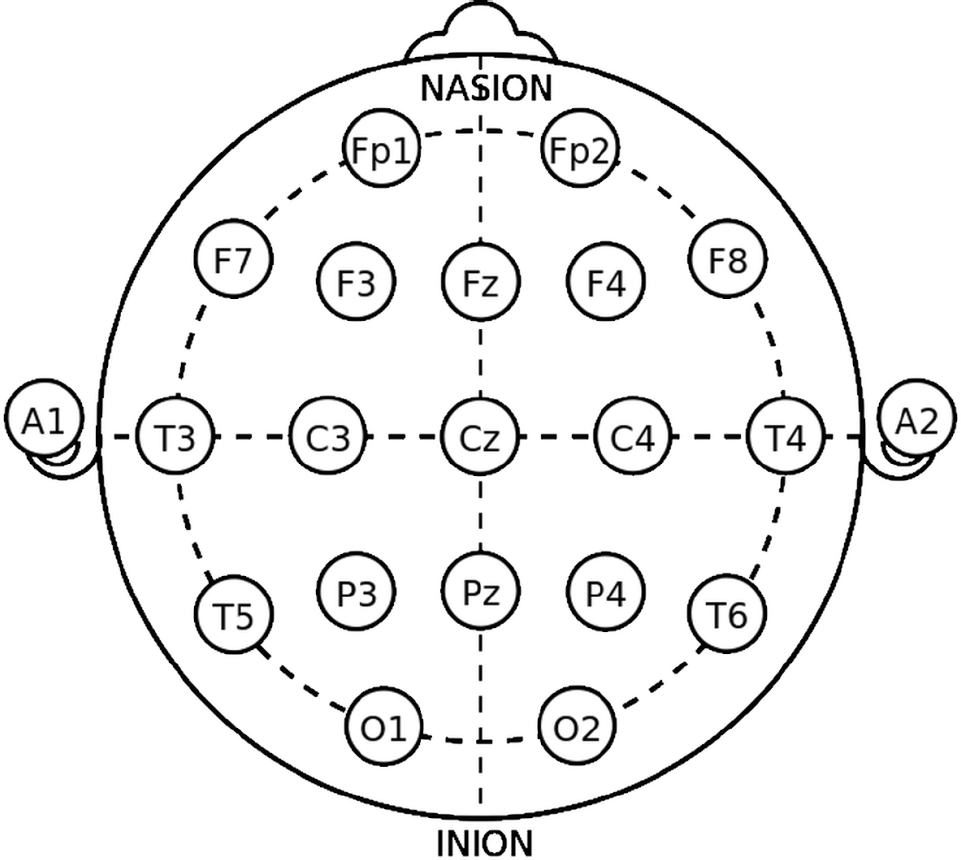
The 10-20 International system of EEG electrode placement.

**Figure 2:**
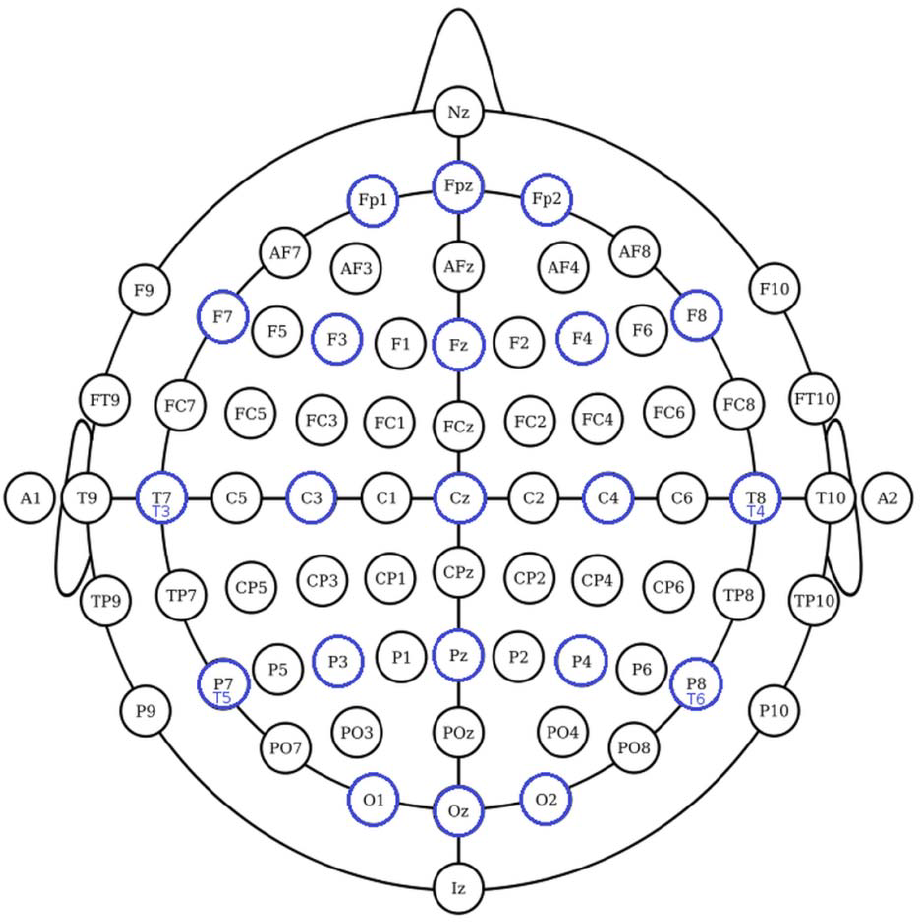
The 10-10 International system of EEG electrode placement. Blue circles represents the location of 10-20 EEG electrodes. Nodes T8, T7, P8 and P7 from 10-10 EEG placement are equivalent to nodes T4, T3, T6, T5 from 10-20 EEG placement.

Functional connectivity MRI (fcMRI) measures the intrinsic functional correlations between brain regions (Mueller et al., 2013; Van Dijk et al., 2010). The method is sensitive to the coupling of both distributed as well as adjacent areas (Biswal et al., 1995; Yeo et al., 2011). It is believed that low-frequency fluctuations observed in the BOLD signals reflect the spontaneous neural activity and that the synchronized fluctuations in distinct brain regions, therefore, point to functional connections between them. The first functional connectivity network found was related to the human motor cortex (Biswal et al., 1995). Later different functional connectivity networks have been found, and these networks change in patients with multiple pathologies (neurological, psychiatric). This renders fcMRI an interesting technique to further our understanding of brain function in health and disease. There are several methods for processing the BOLD signal to obtain connectivity networks: ICA (Independent Component Analysis) (Beckmann et al., 2005; Calhoun et al., 2006), seed-based method using correlation between BOLD time series (Biswal et al., 1995; Margulies et al., 2007; Van Dijk et al., 2010), ALFF (Amplitude of Low-Frequency Fluctuations) (Zang et al., 2007).

Yeo et al. (Yeo et al., 2011), using resting-state data of 1000 healthy individuals, with 1175 ROIs on the cortex, the correlation between the fMRI time series of each ROI, and a clustering algorithm demonstrated the existence of seven main functional networks and a finer solution for 17 functional networks. The seven functional networks are visual, somatomotor, dorsal attention, ventral attention, limbic, frontoparietal, and default mode networks (Figure 3) (Yeo et al., 2011). The networks found by Yeo et al. (Yeo et al., 2011) have shown to be valid across multiple subjects and robust to changes in the data processing. The existence of these networks provides a key motivation for our research.

**Figure 3:**
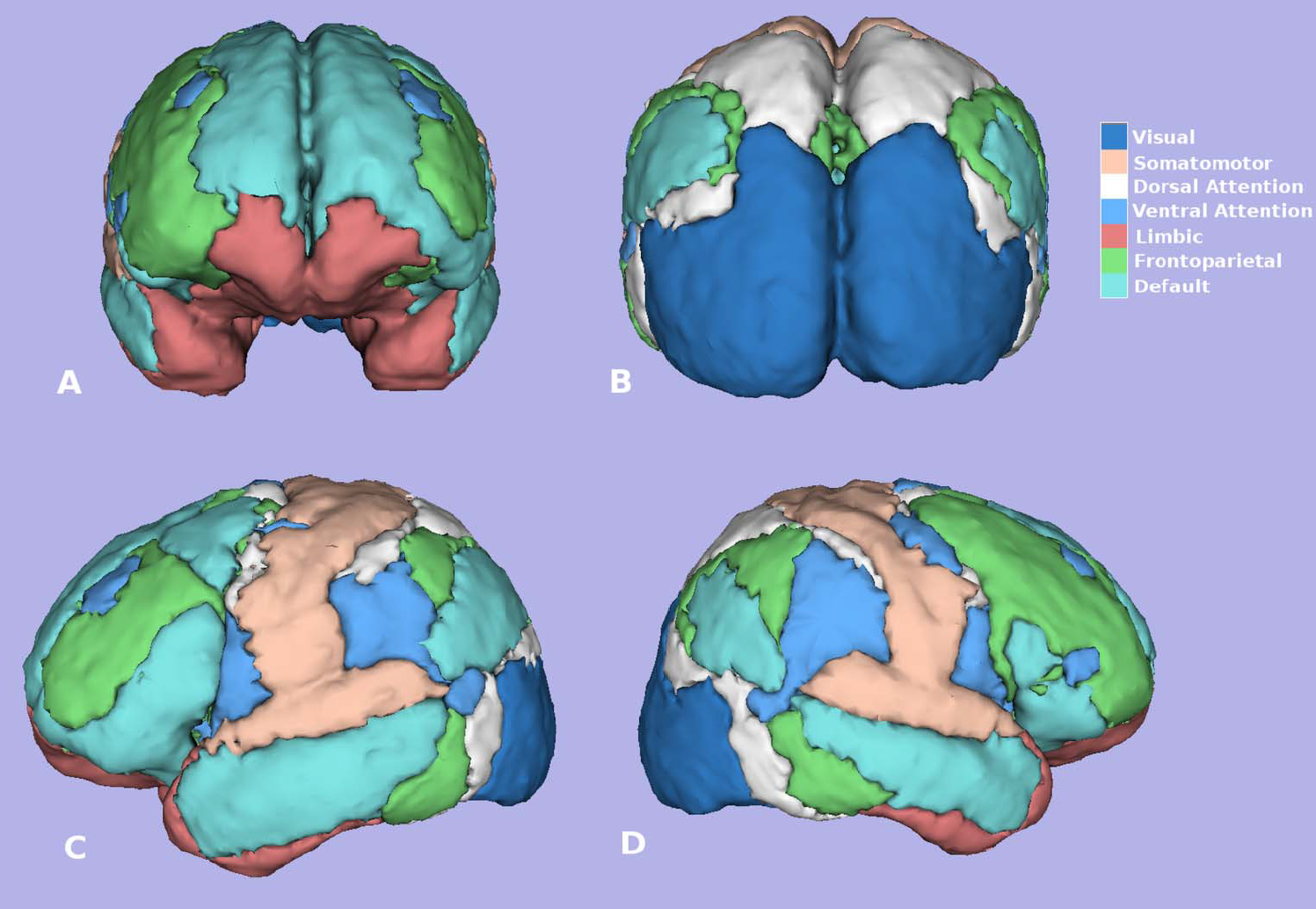
Seven Yeo networks. (Yeo et al, 2011). Figure created using software 3D-Slicer.

One difference between fMRI and EEG is that fMRI has an excellent spatial resolution and a low temporal resolution (seconds), while EEG has a high temporal resolution (milliseconds) with spatial limitations. Simultaneous acquisition of fMRI and EEG could be a solution for the temporal limitations of fMRI and spatial limitations of EEG by combining their features (Ullsperger et al., 2010; Zang et al., 2007). Many authors show different acquisition techniques and data analysis for simultaneous EEG and fMRI (Han et al., 2014; Duyn, 2012; Lazeyras et al., 2001; Menon et al., 2005; Ullsperger et al., 2010). Other difference between both techniques are: EEG measures directly the electrical activity of the brain and fMRI indirectly by measuring changes in blood flow. EEG can be acquired simultaneously with fMRI (high-temporal-resolution data with high-spatial-resolution respectively) (Han et al., 2014; Duyn, 2012; Lazeyras et al., 2001; Menon et al., 2005; Ullsperger et al., 2010).

The fMRI acquired -either simultaneously or not- with an EEG of the same patient, could be processed in many different ways, for instance, ICA (data-driven) (Beckmann et al., 2005; Calhoun et al., 2006), ALFF (Zang et al., 2007), and correlation analysis (seed-based method) (Biswal et al., 1995; Margulies et al., 2007; Van Dijk et al., 2010). In the latter case, it is better to process rs-fMRI using seeds because it is possible to process it in a similar way to EEG (temporal signals in both cases in same places), therefore making it possible to compare the results of both techniques.

In this technical article, we show the functional connectivity networks obtained using seeds relative to the position of 10-20 and 10-10 EEG electrodes, and the relationship of these networks to the 7 and 17 functional connectivity networks (Yeo et al., 2011).

## 2 Materials and Methods

### Subjects

We processed rs-fMRI scans of 45 right-handed healthy volunteers (18-27 years, 10 male and 35 female, TR = 3000 msec, slices = 47, # timepoints = 119, 3T MRI; Cambridge-Buckner dataset, 1000 Functional Connectomes Project; http://fcon_1000.projects.nitrc.org/).

### fMRI data processing and time series analysis

#### Image Preprocessing

Data processing was performed using the Analysis of Functional NeuroImages software (AFNI; http://afni.nimh.nih.gov/afni; Cox, 1996; Cox, 1997) and fMRIB Software Library (FSL; http://fsl.fmrib.ox.ac.uk/fsl/fslwiki; Jenkinson et al., 2012; Smith et al., 2004). Image preprocessing involved the following steps: discarding the first 4 EPI volumes from each resting state scan to allow for signal equilibration; slice-time correction for interleaved acquisitions; 3-D motion correction with Fourier interpolation (volumes in which head motions caused a displacement of more than 2 mm in the x, y, or z-direction, or in which 2° of any angular motion was observed during the course of the scan, were excluded); despiking (detection and removal of extreme time series outliers); spatial smoothing using a 6mm FWHM Gaussian kernel; mean-based intensity normalization of all volumes by the same factor; temporal bandpass filtering (0.009 - 0.1 Hz); and linear and quadratic detrending.

FSL *FLIRT* was used for linear registration of the high-resolution structural images to the MNI152 template (Jenkinson et al., 2001; Jenkinson et al., 2002). This transformation was then refined using *FNIRT* non-linear registration (Andersson et al., 2007a; Andersson et al., 2007b). Linear registration of each participant’s functional time series to the high-resolution structural image was performed using *FLIRT*. This functional-to-anatomical co-registration was improved by intermediate registration to a low-resolution image and b0 unwarping.

#### Nuisance signal regression

To control the effects of motion and physiological processes (related to cardiac and respiratory fluctuations), we regressed each participant’s 4-D pre-processed volume on nine predictors that modeled nuisance signals from white matter, cerebrospinal fluid, the global signal, and six motion parameters. Each participant’s resultant 4-D residuals volume was spatially normalized by applying the previously computed transformation to MNI152 standard space, with 2mm^3^ resolution.

#### Seed Regions of Interest

MNI coordinates corresponding to the 65 electrodes of the 10-10 EEG system were obtained. The algorithm to determine the coordinates related to 10-10 EEG seeds is:

1. Position spheres on the head using MNI coordinates published by Koessler (Koessler et al., 2009; Oostenveld et al., 2001). (Figure 4A)
2. Determine the center of mass of the brain using 3DsMax software (Autodesk Inc., www.autodesk.com) and mesh created with MNI152_T1_2mm_brain_mask_nii.gz (fMRIB Software Library FSL; http://fsl.fmrib.ox.ac.uk/fsl/fslwiki; Jenkinson et al., 2012; Smith et al., 2004). Figure 4B.
3. Calculate the magnitude and direction of the vector connecting the center of mass of the brain and the position of each electrode (Figure 4C).
4. The magnitude of the vector is modified until the electrode sphere (4 mm radius) is completely within the brain by keeping α, β, γ angles unmodified (Figure 4D).

**Figure 4:**
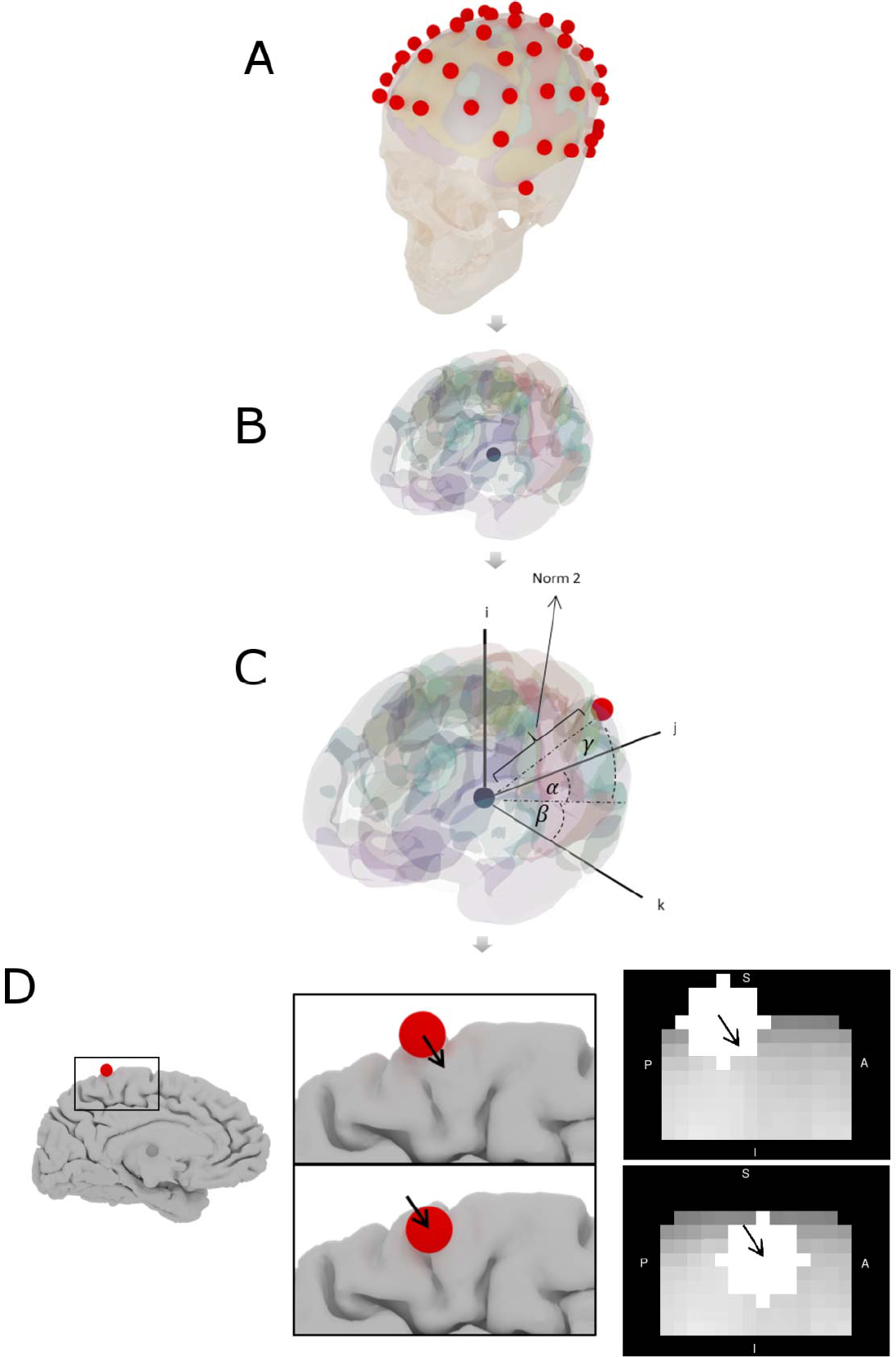
Algorithm to determine 10-10 EEG seeds coordinates. (**A**) Spheres in the head using Koessler coordinates (Oostenveld et al., 2001; Koessler et al., 2009), (**B**) center of mass of the brain, (**C**) magnitude and direction of the vector connecting the center of mass and the position of each electrode, and (**D**) magnitude of the vector is modified until the sphere is within the brain.

In Supplementary Table 1, the coordinates, brain lobes, hemisphere, brain region, Brodmann area and EEG electrode name of each 10-10 EEG ROIs are included. In Supplementary Table 2, the same data is included, but for the ROIs equivalent to the 10-20 EEG electrode system. In Figure 5, the 21 spherical ROIs are shown (10-20 EEG electrode system), and in Figure 6, the 65 spherical ROIs are shown over a brain surface (10-10 EEG electrode system). Figures 5 and 6 were created using BrainNetViewer software (http://www.nitrc.org/projects/bnv; Xia et al., 2013).

**Figure 5:**
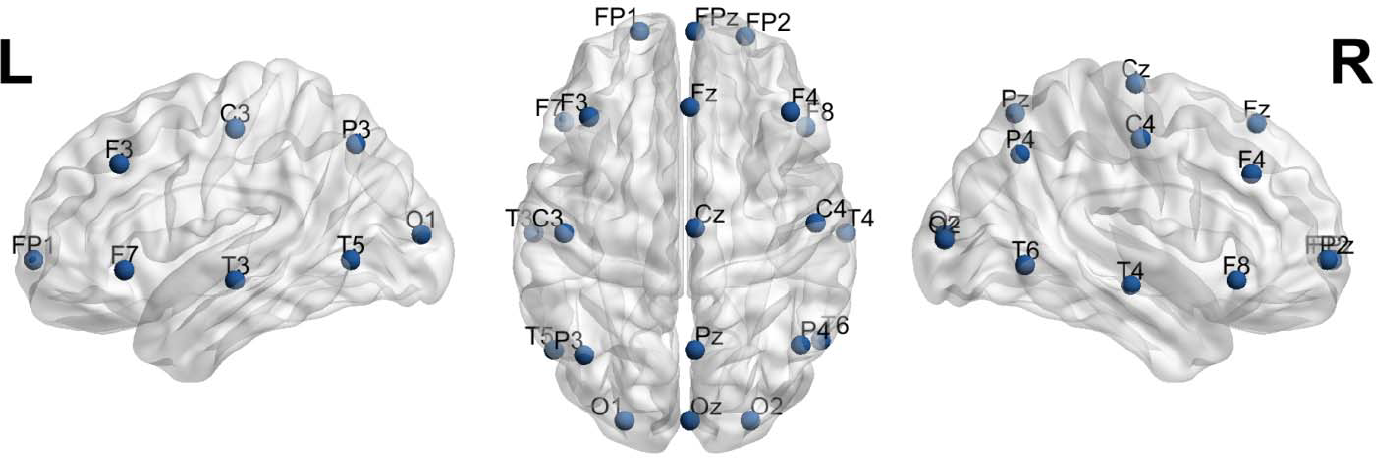
21 spherical ROIs over a brain surface (10-20 EEG electrode system). Figure created using BrainNetViewer software (http://www.nitrc.org/projects/bnv; Xia et al., 2013).

**Figure 6:**
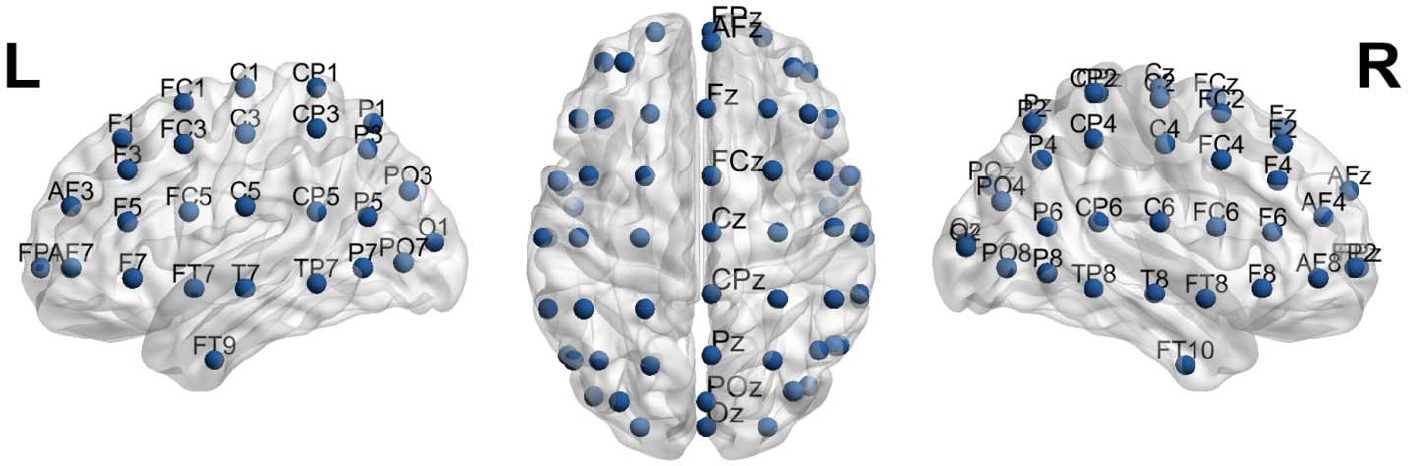
65 spherical ROIs over a brain surface (10-10 EEG electrode system). Figure created using BrainNetViewer software (http://www.nitrc.org/projects/bnv; Xia et al., 2013).

#### Subject-level Resting State Functional Connectivity (RSFC) analysis

For each participant, the representative time series for each 10-10 EEG (seed) ROI was extracted from their 4D residuals volume in standard space by averaging the time-series across all voxels within the ROI. We then calculated the correlation between each seed ROI time series. The resultant participant-level correlation maps were transformed via Fisher-z to Z-value maps and transformed into MNI152 2mm standard space for group-level analyses.

A similar analysis such as the previous one has been carried out using the representative time series for each 10-20 EEG (seed) ROI.

#### Group-level RSFC analysis

For each seed, group-level analyses were carried out using a random-effects ordinary least squares model. Whole-brain correction for multiple comparisons was performed using Gaussian Random Field Theory (min Z>2.3; cluster significance: p<0.05, corrected). This group-level analysis produced threshold z-score maps of activity associated with each 10-10 EEG and 10-20 EEG seed.

### Functional Connectivity networks comparison

To compare the similarity of functional connectivity mapping obtained using each 10-20 EEG electrode seeds and relative to the seven Yeo networks (Yeo et al., 2011), we compute the Sørensen-Dice similarity coefficient (Dice, 1945; Sørensen, 1948). Sørensen-Dice coefficient (also known as Dice index, Sørensen index, or F1 score) is a statistical index commonly used for comparing the similarity of two different samples and their outputs are in the interval between 0 and 1. The Sørensen coefficient was computed using the equation:

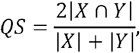

where |*X*|, and |*Y*| is the cardinality of sets X and Y respectively. In our case, is the quantity of voxels of the functional connectivity networks obtained with each 10-20 EEG related seeds, and |*Y*| corresponds to the quantity of voxels of each of the seven Yeo networks (Yeo et al., 2011; https://surfer.nmr.mgh.harvard.edu/fswiki/CorticalParcellation_Yeo2011). In both cases we used MNI152 2mm normalized functional connectivity networks.

## 3 Results

We computed the correlation matrix of each 10-20 EEG electrode seed for each healthy volunteer. The mean matrix of the 45 correlation matrices of each healthy volunteer is depicted in Figure 7A.

**Figure 7:**
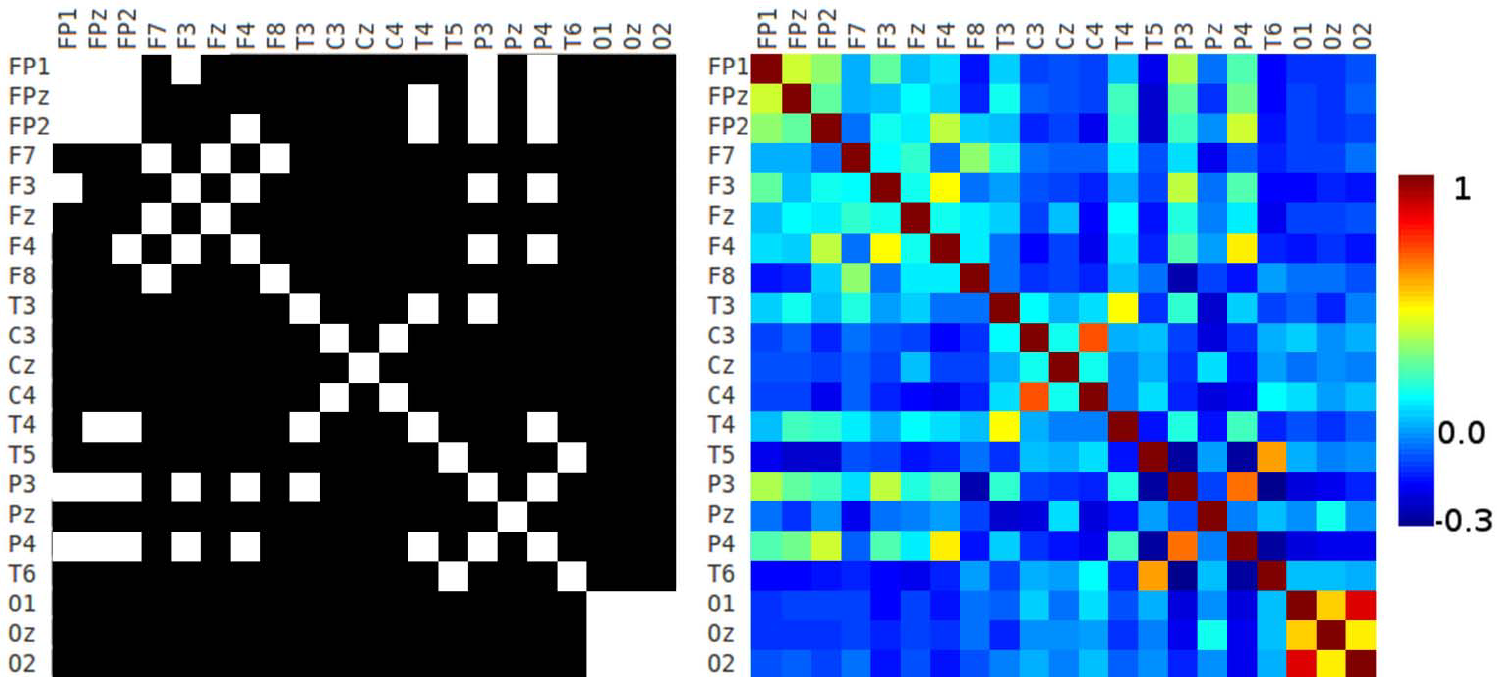
10-20 correlation matrix. (**A**) Correlation matrix of each 10-20 EEG electrode seed and (**B**) adjacency matrix generated using a threshold of 0.2.

The adjacency matrix is a square matrix which is used as a way of representing binary relations, and it shows more clearly the pairs of seeds that have correlation coefficient values greater than a given threshold. Figure 7B shows the adjacency matrix created by applying a threshold of 0.2 to the correlation matrix (Figure 7A). There is strong connectivity between frontopolar and some frontal 10-20 EEG seeds. In Figures 7A and 7B appear contralateral strong connectivity between frontal, central, temporal, parietal, and occipital seeds. Also, P3 and P4 have strong connectivity with frontopolar and some frontal seeds. Occipital seeds have high connectivity between them.

Figure 8A shows the correlation matrix of the 10-10 EEG electrode system seeds, and Figure 8B shows the adjacency matrix with a threshold of 0.2 (mean value of positive values of the correlation matrix). These results are similar to Figure 6, but with higher resolution (due to the fact that 10-10 EEG has more seeds than 10-20 EEG). Frontopolar, anterior-frontal, frontal, parieto-occipital, and occipital seeds have ipsilateral and contralateral strong connectivity. Some frontocentral, central, centroparietal and parietal seeds have contralateral high connectivity (Figures 8A and 8B).

**Figure 8:**
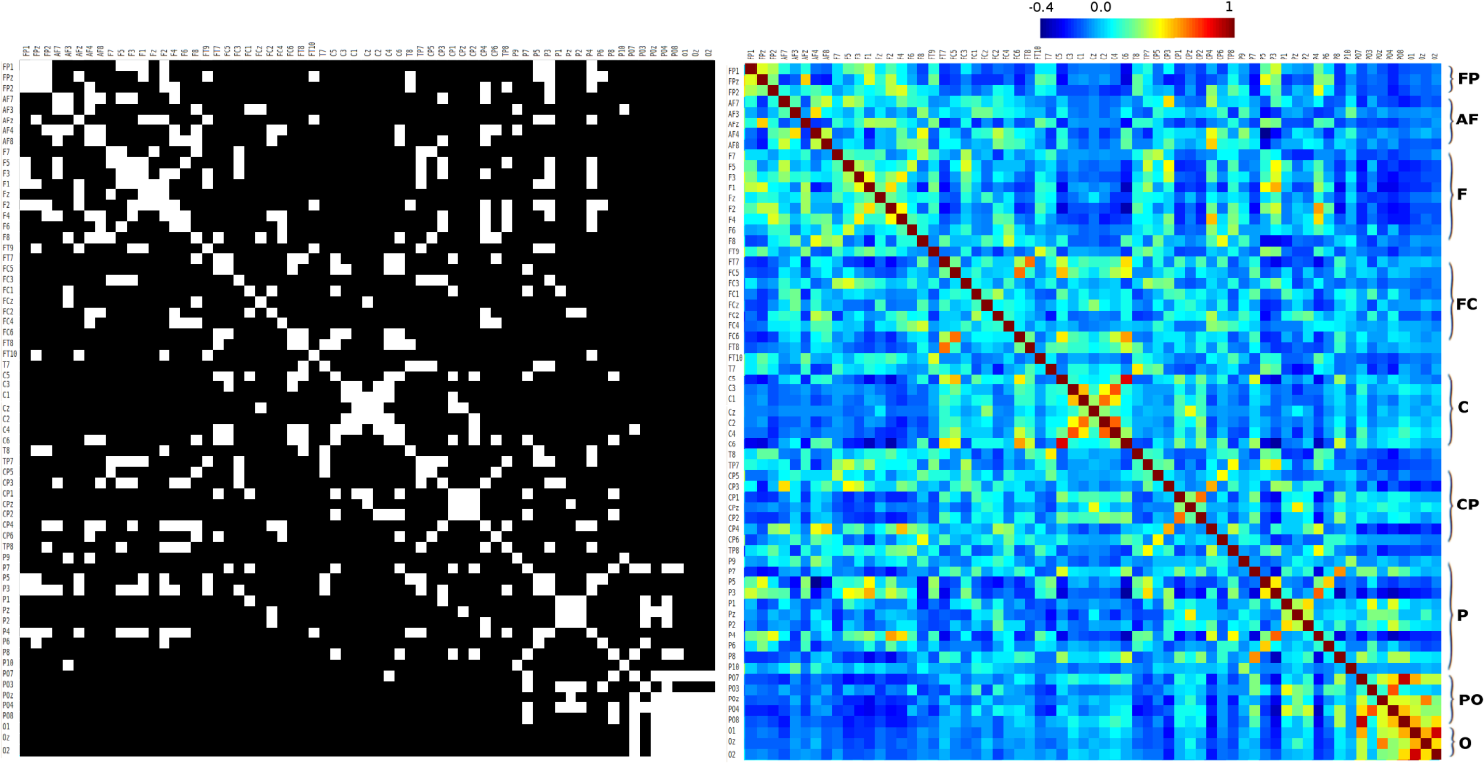
10-10 correlation matrix. (**A**) correlation matrix of 10-10 EEG electrode seed system, and (**B**) adjacency matrix generated using a threshold of 0.2. **FP**: Fronto-polar, **AF**: Anterior-frontal, **F**: Frontal, **FC**: Fronto-central, **C**: Central, **CP**: Centro-parietal, **P**: Parietal, **PO**: Parieto-occipital, **O**: Occipital.

We computed Sørensen-Dice coefficient. In Supplementary Table 3, we show the Dice similarity index related to Visual, Somatomotor, Dorsal Attention, Ventral Attention, Limbic, Frontoparietal, and Default mode networks. Figure 9 depicts the Dice indices as a bar chart in each 10-20 EEG seed. Broadly, Dice index does not have large values (less than 0.68), and higher-value Sørensen-Dice coefficients are in occipital seeds. Sørensen-Dice coefficient shows laterality for some Yeo networks.

**Figure 9:**
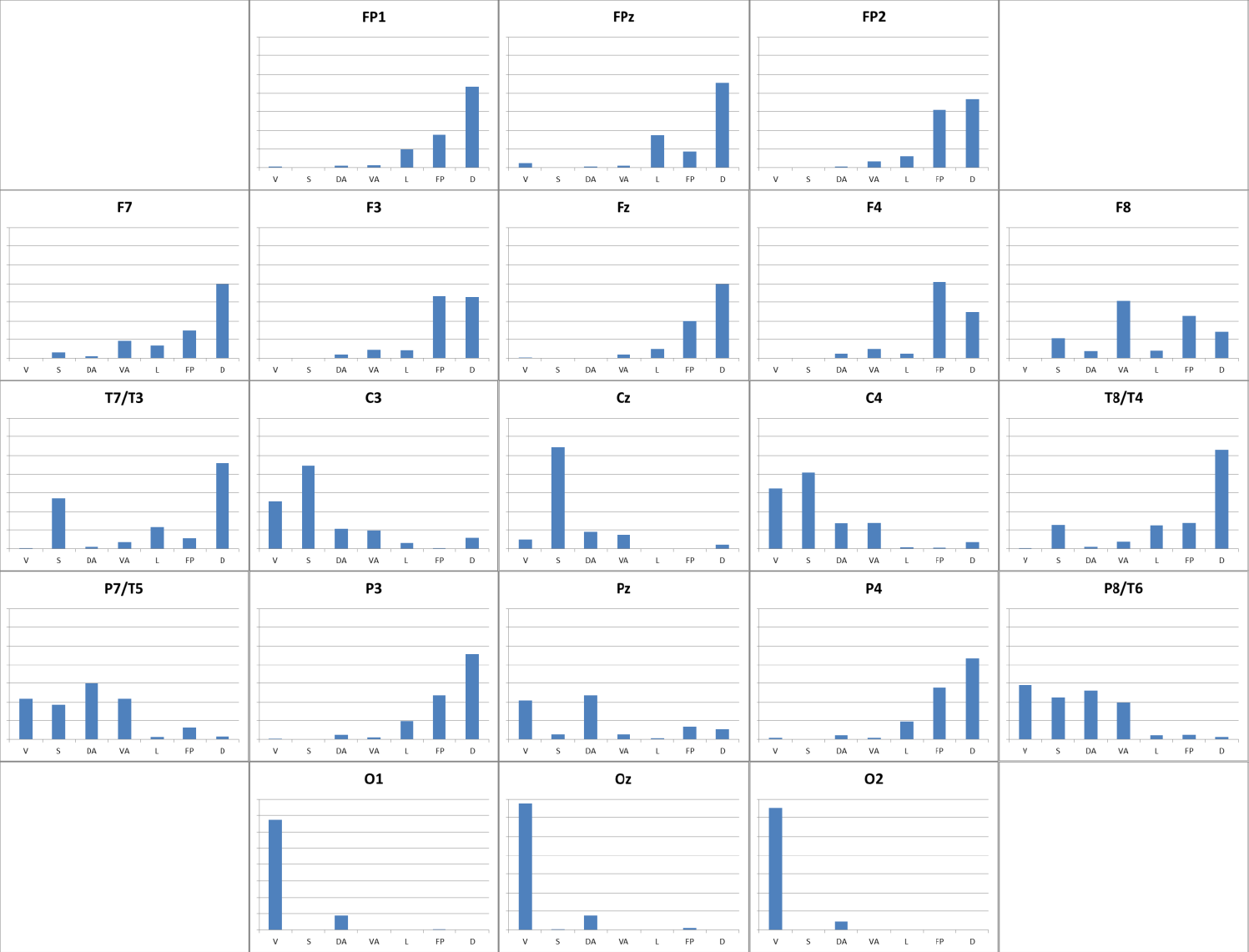
Similarity of 10-20 EEG seeds relative to Yeo networks. The similarity of functional connectivity mapping obtained using each 10-20 EEG seeds relative to seven Yeo networks (Yeo et al., 2011). The figure shows a bar chart for each EEG seed with Sørensen-Dice similarity coefficient (Sørensen, 1948; Dice, 1945) (see Material and Methods section). In each bar chart X-axis shows Yeo networks (**V**: Visual, **S**: Somatomotor, **DA**: Dorsal attention, **VA**: Ventral attention, **L**: Limbic, **FP**: Frontoparietal, **D**: Default mode network; columns from left to right), and Y-axis shows Sørensen-Dice similarity coefficient (0.0 to 0.8 values).

Figures 10 and 11 show 3D functional connectivity surface maps for each 10-20 EEG electrode seed. The maps were created with the 45 previously mentioned rs-fMRI images (Cambridge-Buckner dataset, 1000 Functional Connectomes Project).

**Figure 10:**
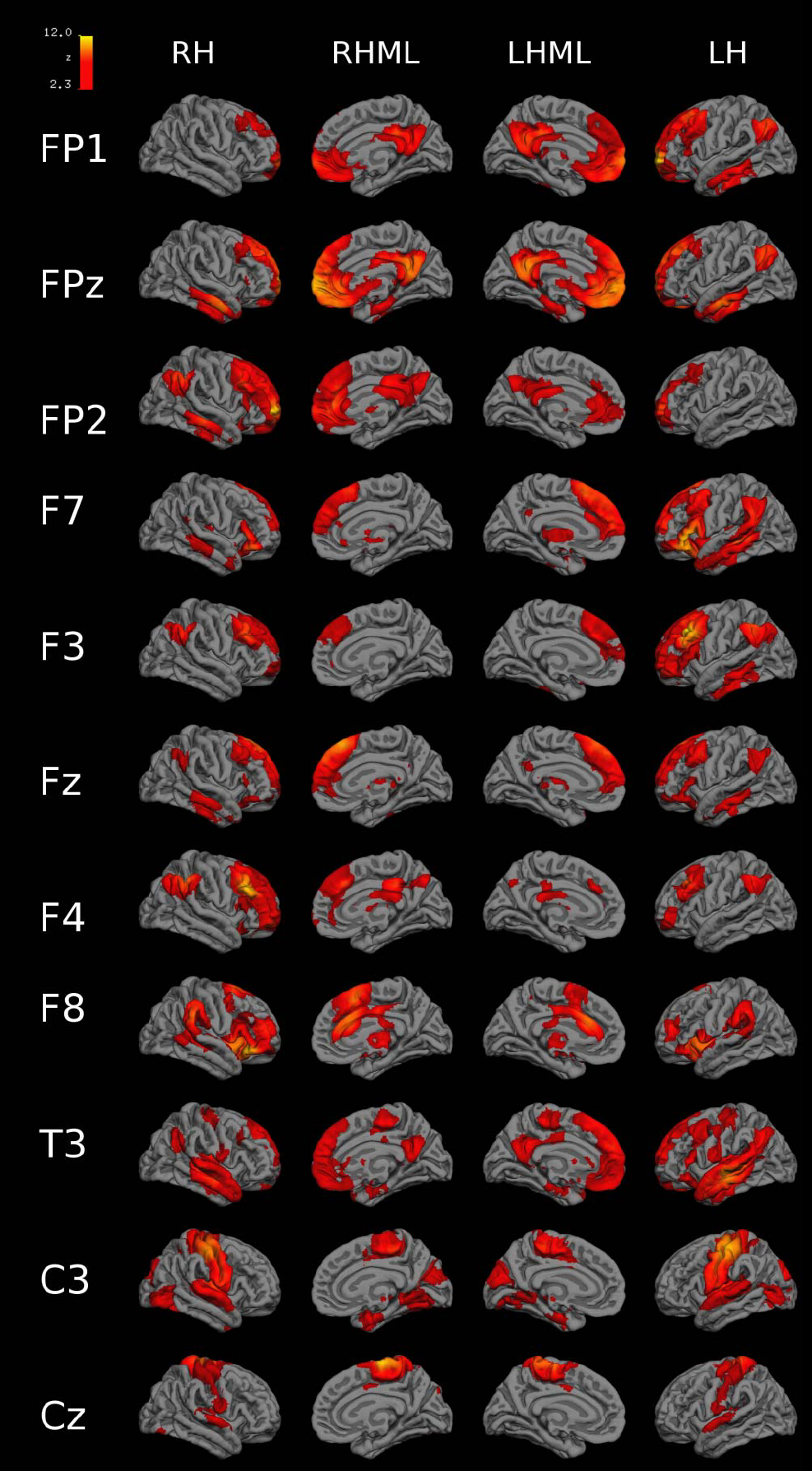
3D functional connectivity surface maps (FP1-Cz seeds). 3D connectivity images for each 10-20 EEG electrode-seed (FP1 to Cz) created with 45 rs-fMRI images previously mentioned in Material and Methods section (Cambridge-Buckner dataset, 1000 Functional Connectomes Project).

**Figure 11:**
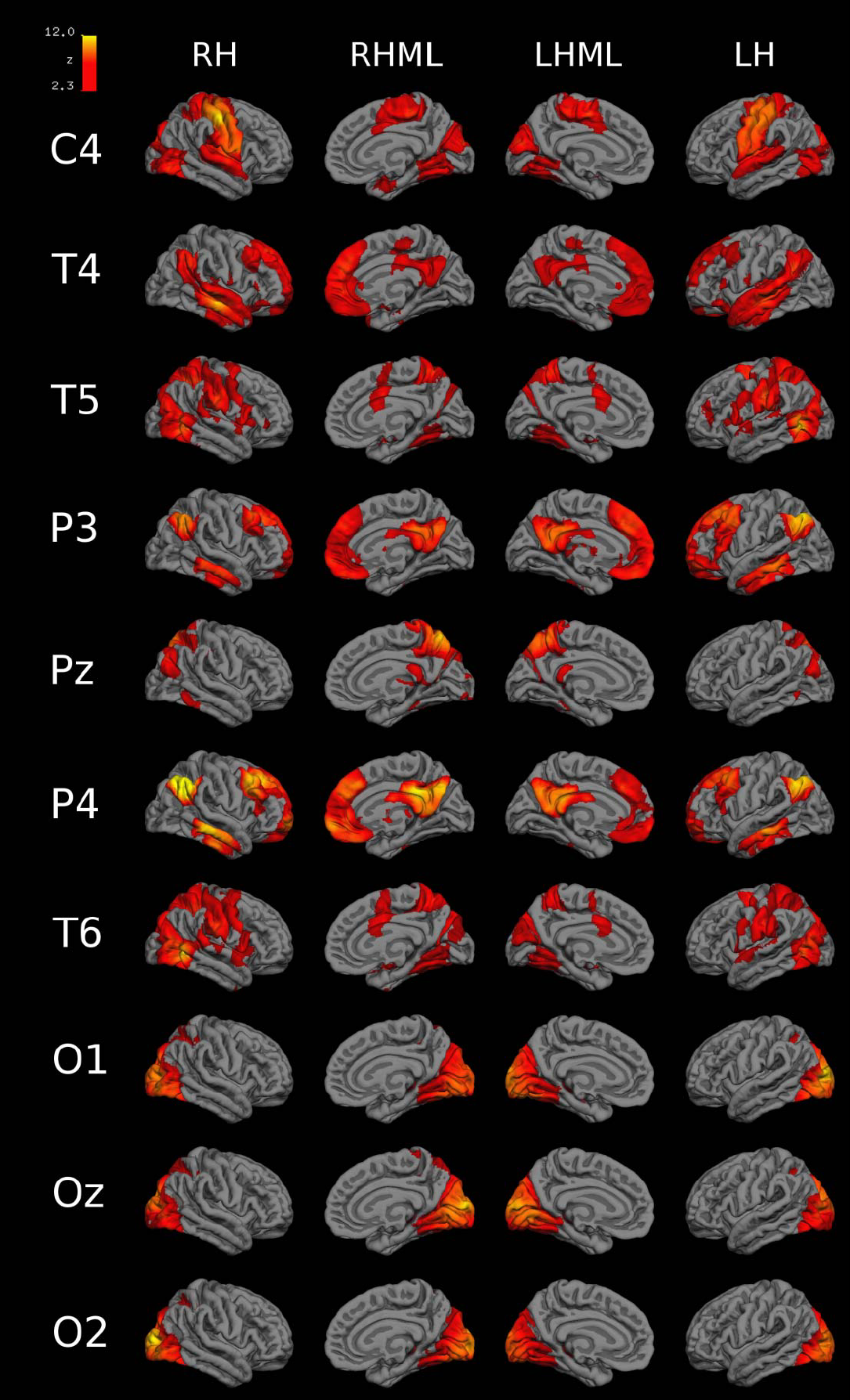
3D functional connectivity surface maps (C4-O2 seeds). 3D connectivity images for each 10-20 EEG electrode-seed (C4 to O2) created with 45 rs-fMRI images previously mentioned in Material and Methods section (Cambridge-Buckner dataset, 1000 Functional Connectomes Project). Associative areas have high long-range connections that are related to greater functional variability. Apparently, distant connectivity is necessary for prominent functional variability, and this is not observed in species with smaller brains. On the other hand, association areas are more recent in the evolution of the human brain development, whose greater functional variability could also be explained by its later maturation during development, which would make these more vulnerable to external postnatal influences and less dependent on genetic factors (Mueller et al., 2013).

In Figures 10 and 11 FP1, FP2 seeds showed connectivity with bilateral medial prefrontal cortex (MPF), bilateral posterior cingulate cortex (PCC), ipsilateral lateral parietal cortex (LP), and ipsilateral inferior temporal cortex (IT). FPz connects with bilateral MPF, bilateral PCC, left hemisphere LP, bilateral IT and bilateral parahippocampal cortex. F7 connects with bilateral thalamus, ventrolateral prefrontal cortex (VLPFC), IT, and MPF, and ipsilateral LP. F8 has connectivity with bilateral inferior parietal gyrus, anterior dorsolateral PFC (aDLPFC), ipsilateral MPF, contralateral dorsal anterior cingulate cortex (dACC), and insula. F3 connects with bilateral MPF, and LP, ipsilateral aDLPFC, and IT. F4 connects with ipsilateral aDLPFC, PCC, and bilateral LP. Fz has functional connectivity with bilateral MPF, IT, and LP. T3 and T4 have connectivity with bilateral LP, PCC, IT, middle temporal gyrus, and superior temporal gyrus, ipsilateral VLPFC and precentral gyrus (only with T3). C3 and C4 connect with bilateral pre/post central gyrus (PCG), posterior occipital cortex (pOCC), cuneus, superior temporal gyrus, parahippocampal gyrus, and fusiform gyrus. Cz has connectivity with bilateral PCG and superior temporal gyrus. T5 and T6 have connectivity with bilateral pOCC, IPS, PCG, superior frontal gyrus, superior temporal gyrus, lateral occipital-temporal gyrus, and aDLPFC (only in T6). P3 and P4 connect to bilateral LP, MPF, IT, middle temporal gyrus. Pz connects with right hemisphere LP, bilateral intraparietal sulcus (IPS), and PCC. O1 connect with bilateral precuneus and hippocampus, and ipsilateral cingulate gyrus. O2, with bilateral precuneus, hippocampus, and cingulate gyrus. Oz has a connection with bilateral precuneus and hippocampus and left hemisphere cingulate gyrus.

## 4 Discussion

Epilepsy and other neurological diseases are diagnosed with EEG. In this paper, we show rs-fMRI results and analysis processed in a similar way as EEG.

In most clinical cases, the 10-20 EEG electrodes scheme is used adding some 10-10 EEG electrodes, and, therefore, functional connectivity was processed in a similar way as EEG, i.e. using 10-20 and 10-10 EEG related seeds.

On the basis of the correlation and 10-20 adjacency matrixes (Figures 7A and 7B), it can be suggested the following:

Broadly, there is a high correlation between contralateral seeds (for instance FP1-FP2, F3-F4, F7-F8, C3-C4, T3-T4, P3-P4, and T5-T6). Also, there is a high correlation between ipsilateral seeds (FP2-F4, FP1-F3, FP2-T4), high correlation between frontopolar, frontal and anterior temporal seeds with parietal seeds (FP1-P3, FP2-P3, FPz-P3, F3-P3, F3-P4, F4-P3, F4-P4 and T3-P3), and high correlation between occipital seeds: O1-O2, Oz-O1, Oz-O2.

The 10-10 correlation matrix (Figure 8A), suggests that:

There is a high connectivity between frontal, frontopolar, anterior-frontal, and frontocentral EEG seeds, between ipsilateral and contralateral seeds (adjacency matrix with a threshold of 0.2 in Figure 8B). There is a high contralateral connectivity between centroparietal, and parietal EEG seeds.

There is high connectivity between the occipital EEG seeds (between ipsilateral and contralateral seeds; Figure 8B).

The occipital EEG seeds (O1, O2, Oz) only have connectivity with PO7 (Figure 8B).

### Analysis for each functional connectivity network

In eleven EEG seeds (FP1, FP2, FPz, F3, F4, F7, Fz, T3, T4, P3, P4), the Sørensen-Dice coefficient (Figure 9) of the **Default Mode Network** is greater than 0.2, and the left hemisphere EEG seeds have a higher Sørensen-Dice coefficient than those of the right side (left lateralized functional connectivity network). Our results are consistents with (Agcaoglu et al., 2015) that specify the same lateralization of the Default mode network for young people, and similar left lateralization conclusions reports (Nielsen et al., 2013) and (Swanson et al., 2011) for healthy people.

The **Frontoparietal Network** is right-lateralized because it has a higher percentage of functional connectivity voxels in even numbered EEG seeds than in odd numbered seeds. The Frontoparietal Network coexists with the default mode network (more than 35% of maximum value Sørensen-Dice index; 0.143 and 0.186 respectively) in multiple frontal seeds (Fp1, Fp2; BA 10, F3, F4; BA 9, FZ; BA 8), anterior temporals (F7, BA 47) and parietal seeds (P3, P4, BA 39).

**Ventral Attention Network:** the Sørensen-Dice coefficient has the highest value in F8 seed (BA 47; 0.306). Other high values of Sørensen-Dice coefficient are in T5, T6, C4 (0.218, 0.199, and 0.138 respectively).

C3 (BA 3), C4 (BA 3), T5 (BA 19), T6 (BA 19), and Pz (BA 7), have a Dice index greater than 0.105 (35% of maximum value of Dice index) for the **Dorsal Attention Network**. This network is not lateralized (4.3 % difference between left and right mean Sørensen-Dice coefficient).

**Limbic Functional Network** is also left-lateralized because in general, odd numbered EEG seeds have higher Dice index than the even numbered ones. And because, the percent difference between left and right mean Sørensen-Dice coefficient is more than 20%.

The **Somatomotor functional network** has the highest Sørensen-Dice index in Cz, (BA 6), C3 (BA 3) and C4 (BA 3) EEG seeds (0.544, 0.447, and 0.410 respectively). In general, we could conclude that it is an equally lateralized network (less than 8% difference between both hemispheres of Sørensen-Dice coefficient).

The **Visual Functional Network** has a Dice index greater than 0.600 in O1 (BA 19; Associative visual cortex V3, V4, V5), 02 (BA 18; secondary visual cortex V2), and Oz seeds. In C4 (BA 3), T6 (BA 19), and C3 (BA 3) the Visual Network has a Sørensen-Dice coefficient more than 0.25. In the connectivity images for O1, O2, and Oz appears to have connectivity only with occipital and parietal regions (Figure 8, Figure 11, Supplementary Table 3). Using Dice index (10% difference) we could specify that Visual functional network is right lateralized as specified (Agcaoglu et al., 2015).

### General Analysis

The fact that more than one network coexists in a single seed (see bar charts in Figure 9) can be explained because the brain is and works as a network of interconnections (or graph-connected) where the existing networks are not independent and must be connected to function as an integrated organ. If we assume that networks are independent, different functional areas such as memory, visual or, motor could not be inter-related, contradicting the fact that the brain works as a network of interconnections.

Seeds located according to 10-20 EEG system in frontopolar, frontal, anterior temporal and parietal regions show high regional connectivity in relation to the default mode and frontoparietal networks. The default mode network predominates in the left hemisphere, probably this being related to a higher representativity of this functional network to the left, whereas in the right hemisphere the frontoparietal networks predominate. The frontoparietal network, located between the default mode network and the dorsal attention network, has a role in goal-directed cognition (Spreng et al., 2013; Vincent et al., 2008), and in the integration of information from the dorsal attention network and the default mode network (Vincent et al., 2008).

Also, frontal contralateral seeds have high connectivity (Figures. 7, 8 and 10). Similar results were published in an ICA-derived EEG functional connectivity study that shows the power spectra of each independent component (Brodmann area 10, Figure 1C in Chen et al., 2013), and other EEG connectivity based paper that specify that contralateral frontal connections are common (61%, Figure 3 in Lacruz et al., 2007).

Central level seeds (C3, C4, and Cz) have high connectivity between each other with significant somatomotor representation (primary motor-sensory area; Figures 10 and 11), and moderate connectivity with temporal posterior region seeds (Figures 10 and 11) coinciding with the dorsal attention network (functional area related to processing of external sensory stimuli), ventral attention network, and not connected with the default mode network region (Buckner et al., 2013).

Seeds in the occipital region (visual area) have high ipsilateral and contralateral connectivity slightly connected to some dorsal attention network regions (precuneus, cingulate gyrus, and hippocampus Figure 10). The primary sensory areas (visual, somatomotor) and the limbic area would be evolutionarily older than other parts of the prosencephalon, which is consistent with a modular-type organization, probably determined phylogenetically. Primary sensory areas participate in simple networks with local networks preferably. The greater the local connectivity is, the less functional variability exists, i.e. the occipital region is a stable connectivity area between the individuals (robust networks) and, therefore, would have no variability with respect to their connections. Instead, areas represented by seeds in associative area regions, mainly prefrontal, temporal and parietal, coinciding with frontoparietal and attention networks, have connectivity above the global mean of the inter-subject variability (Yeo et al., 2013). Fahoum et al. (Fahoum et al., 2012) shows that in posterior quadrant epilepsy there exist only deactivation clusters in bilateral PCC and precuneus, not activations.

Being able to establish normal connectivity models through combination (or fusion) of methods -in this case, electric signals and hemodynamic data-allows us to address the temporal and spatial relationship of the observed connections, and, in the future, to correlate it with connectivity in epilepsy patients (or other brain disease), in order to establish abnormal association networks so as to not only try to refine the detection of seizure origin area using non-invasive methods, but also to predict surgical prognosis, response to drug therapy, or cognitive impairment, inter allia.

The functional connectivity analysis of the rs-fMRI using 10-20 EEG positioned seeds as proposed in this article makes it possible to obtain connectivity maps superimposed with networks described by Yeo (Yeo et al., 2013) and Brodmann areas (Brodmann, 1909), determining comparable connectivity matrices.

Functional connectivity analysis of rs-fMRI using 10-10 or 10-20 EEG positioned seeds helps the physician to understand the connectivity alterations detected by fMRI using classical parameters of EEG localization. EEG is currently the standard method for monitoring or diagnosis of some diseases (Niedermeyer et al., 1993). Also, using the proposed scheme it is possible to perform a correlation analysis between the anatomical layout of EEG surface electrodes and Yeo functional connectivity networks (Rojas et al., 2016; Yeo et al., 2011).

Finally, the objective of our study was to provide a replicable model in which the arrangement of seeds using 10-20 EEG system allows us to combine methods for temporal and spatial location through noninvasive markers, in order to identify abnormal functional networks.

